# *Streptococcus anginosus* Activates the NLRP3 Inflammasome to Promote Inflammatory Responses from Macrophages

**DOI:** 10.1101/2025.03.12.642696

**Authors:** Anika M. Arias, Dakota M. Reinartz, Chloe Sairs, Sangeetha Senthil Kumar, Justin E. Wilson

## Abstract

Chronic inflammation and oral dysbiosis are common features of oral squamous cell carcinoma (OSCC). The commensal streptococci, *S. anginosus,* is increased in oral diseases including OSCC. Our previous work revealed that *S. anginosus* promotes inflammatory responses from macrophage cell lines, however the molecular mechanism by which *S. anginosus* interacts with macrophages to instigate this response remains to be investigated. Here, we expand on our previous findings by investigating the effects of *S. anginosus* infection of primary bone marrow derived macrophages (BMMs) and during *in vivo* infection. We found *S. anginosus* activated primary BMMs, which presented an enlarged cellular area, increased NF-κB activation and downstream inflammatory cytokines TNF⍰, IL-6 and IL-1β at 24 hours post infection. *S. anginosus* viability was dispensable for NF-κB activation, but essential for the induction of downstream inflammatory proteins and cytokines. *S. anginosus* persisted intracellularly within BMMs and induced the expression of inflammasome sensors AIM2, NLRC4 and NLRP3. Further, BMMs lacking the inflammasome adapter protein ASC (*Asc^−/−^)* had significantly diminished IL-1β production compared to wild type BMMs, indicating that *S. anginosus* activated the inflammasome. *S. anginosus* primarily triggered the inflammasome through NLRP3 as *S. anginosus*-infected *Nlrp3^−/−^* BMMs and NLRP3 inhibitor (MCC950)-treated wild type BMMs displayed diminished IL-1β production compared to wild type controls. Lastly, *S. anginosus*-infected *Asc^−/−^* and *Nlrp3^−/−^*mice displayed reduced weight loss compared to C57BL/6 mice. These overall findings indicate that *S. anginosus* replicates within macrophages and promotes a proinflammatory response in part through activation of the NLRP3 inflammasome.

**brief summary sentence:** *S. anginosus* replicates intracellularly within macrophages and is sensed by the NLRP3 inflammasome to promote proinflammatory response.

## 1. Background

Head and neck squamous cell carcinoma (HNSCC) is the seventh most common type of cancer worldwide [1] with 890,000 new cases and 450,000 deaths annually. HNSCC refers to a diverse group of malignancies arising from the anatomic sites involving the oral cavity, pharynx, hypopharynx, larynx, nasal cavity, and salivary glands that compose the upper aerodigestive tract [2]. Oral squamous cell carcinoma (OSCC) is the predominant form of HNSCC and is responsible for about 389,846 new cancer cases with an estimated 188,438 deaths in 2022 [3]. The number of newly diagnosed OSCC cases is predicted to increase by 62% in 2035 [4]. Despite advances in treatment, OSCC has a poor prognosis, with less than a 50% 5-year survival rate in many regions [5]. While tobacco and heavy alcohol use account for 74% of OSCC cases, 2–6% are linked to human papillomavirus (HPV) infection [6]. Notably, 15% of oral cancer cases occur in individuals without a history of tobacco or alcohol use, suggesting the potential role of other emerging risk factors for OSCC, including oral dysbiosis [7]. These challenges have urged research into new risk factors and prognostic markers, with increasing attention on the role of the oral microbiome in oral carcinogenesis [7].

Dysbiosis of the oral microbiome exacerbates disease largely through prolonged inflammatory signaling, although some bacteria are directly linked to other cancer hallmarks. An increase in pathogenic species such as *Fusobacterium nucleatum* and *Porphyromonas gingivalis* is linked to an increase in pro-inflammatory NF-κB signaling, elevated production of pro-inflammatory cytokines TNF⍰ and IL-1β, and an increase in OSCC proliferation and invasion [8],[9],[10]. Additionally, streptococcus species such as *S. mutans* can increase inflammatory cytokines including IL-6 and promote metastasis by decreasing vascular endothelial cadherin [11]. Other Gram-positive streptococcus species are reportedly elevated in OSCC including *S. constellatus*, *S. salivarius*, *S. gordonii* and *S. mitis*, although their mechanism in disease progression remains to be investigated [12]. Similarly, *S. anginosus* is also increased in OSCC [13]. *In vitro*, the *S. anginosus* virulence factor Streptolysin S increases production of HCS-2 cell immediate early genes (IEGs), which are responsible for transcription of early growth factors (EGFs) [14]. Conversely, *S. anginosus* decreases proliferation, migration and invasion of the SCC-15 cell line [15]. While there is a scarcity of studies investigating the role of *S. anginosus* during OSCC *in vivo*, *S. anginosus* promotes proliferation, inhibits apoptosis and induces gastric tumorigenesis in mice [16].

The host innate immune system plays a crucial role in balancing mucosal immunity versus tolerance to bacteria in the oral cavity. Tissue macrophages represent important contributors of innate immunity through their direct interactions with oral microbes and subsequent responses that direct immune activation. Microbial recognition by macrophages is mediated by pattern recognition receptors (PRRs), which include surface PRRs, including most Toll-like receptors (TLRs) and C-Type Lectin receptors (CLRs), and cytosolic PRRs, including NOD-like receptors (NLRs), RIG like receptors (RLRs), and AIM2-like receptors (ALRs). PRRs are able to recognize a variety of pathogen-associated molecular patterns (PAMPs) and damage-associated molecular patterns (DAMPs) to signal an inflammatory response [17]. TLRs recognize many microbial surface ligands and nucleic acids within endosomes [18]. Upon recognition of its ligand, TLRs upregulate proinflammatory signaling pathways such as NF-κB and induce production of proinflammatory cytokines such as TNFα, IL-6 and type I interferons [18]. Additionally, TLRs can upregulate the expression of intracellular PRRs such as NLRs/ALRs and immature cytokines such as pro-IL-1β and pro-IL-18 [19]. Several intracellular sensors, including NLRP3 and AIM2, form an inflammasome by binding with the adaptor protein ASC and the inactive enzyme pro-caspase-1. Inflammasomes can be activated by a variety of PAMPS and DAMPS (e.g., extracellular ATP, flagellin and lipoteichoic acid), depending on the NLR sensor, while the AIM2 inflammasome is triggered by double-stranded DNA [20], [21]. Inflammasome assembly leads to pro-caspase-1 cleavage into its active form, capasase-1, which subsequently cleaves pro-IL-1β, pro-IL-18 and Gasdermin D. Mature IL-1β and IL-18 are then secreted through pores formed by Gasdermin D to initiate inflammatory responses. During severe infections, the inflammasome can also mediate a Caspase-1-dependent form of inflammatory cell death called pyroptosis [20].

A variety of ligands from differing streptococcus species can both upregulate the expression of and activate TLRs, NLRs and other intracellular sensors. Components of the cell wall of *S. pneumoniae* upregulates numerous TLRs including TLR1, 2, 6 and 10 [22]. Additionally, pilus components of *S. pyogenes*, which are necessary for bacterial adhesion, are recognized by TLR2 [23]. Several intracellular sensors are crucial to innate control of streptococcus species found in the oral cavity as well. NLRP3 recognizes *S. pneumonia* virulence factors pneumolysin and bacterial-derived hydrogen peroxide [24]. *S. pneumoniae* and *S. gordonii* are recognized by NLRP6, although the microbial ligand sensed by NLRP6 was not identified in these studies [25], [26]. However, NLRP6 recognizes lipoteichoic acid found on other Gram-positive bacteria, including *Listeria monocytogenes* [27]. Additionally, AIM2 plays a crucial role in recognizing and mediating an inflammatory response against *S. pneumoniae* infection [28], thus highlighting the coordination of multiple inflammasome sensors during streptococcus recognition. Other intracellular sensors that recognize streptococcus species include NLRC4, which traditionally senses flagellin and type III secretion system components, although it also recognizes bacteria without flagellin and host-derived DAMPs [29] [30] [31]. It is unclear which intracellular sensors recognize *S. anginosus* infection to promote inflammation.

Our previous work using the RAW264.7 macrophage cell line indicates that infection by *S. anginosus* induces robust production of inflammatory cytokines TNF⍰, IL-1β, and IL-6, induction of inflammatory mediators including active NF-κB, iNOS, and COX2, and upregulation of the intracellular sensor NLRP3 [32]. Here, we demonstrate that *S. anginosus* induces a similar proinflammatory protein and cytokine profile in primary bone marrow derived macrophages (BMMs), including IL-1β production. Moreover, *S. anginosus* replicates within resting BMMs, and heat-inactivated *S. anginosus* fails to induce IL-1β, indicating bacterial viability is crucial to mounting a fulminate inflammatory response. Additionally, *S. anginosus* upregulates multiple intracellular sensors including NLRP3, NLRC4 and AIM2. Genetic ablation or pharmacological inhibition of NLRP3 in *S. anginosus*-infected BMMs lead to reduced IL-1β production. Thus, *S. anginosus* recognition and downstream IL-1β production is partially mediated by the NLRP3 inflammasome.

## 2. Materials and Methods

### 2.1 Bacteria

*Streptococcus anginosus* was purchased from the American Type Culture Collection (ATCC 33397) and maintained in a 50% glycerol stock solution at −80°C until use. For all described experiments, *S. anginosus* was grown on blood agar plates containing Tryptic Soy Agar (22091-500G Millipore) and 20% defibrinated donor sheep blood (610-500 Quad Five) at 37°C for 18 hrs. On the day of infection, a loop of plated *S. anginosus* was inoculated into Brain Heart Infusion Broth (53286-500G Millipore) and grown to an OD 600 of 0.1 (5.4×10^7^ CFUs/mL) at 37°C [32].

### 2.2 Mice

All animal protocols were approved by the Institutional Animal Care and Use Committee (IACUC) of the University of Arizona in accordance with the US National Institutes of Health Guide for the Care and Use of Laboratory Animals. Animal numbers were empirically determined to optimize numbers necessary for statistical significance based on our previous reports utilizing these disease models while adhering to IACUC polices on maximum numbers of animals used in research. Animals had access to water and food ad libitum. All experiments were performed under specific pathogen-free conditions using 6- to 8-week-old mice. Wild-type (WT) C57BL/6 mice and *Nlrp3^−/−^* mice were originally obtained from the Jackson Laboratories (Bar Harbor, ME) and maintained at University of Arizona Health Sciences Animal Facility. *Asc^−/−^* mice were generously provided by Dr. Vishva Dixit (Genetech) and housed at the University of Arizona [33]. *Nlrc4^−/−^* mouse bones were generously provided by Dr. Isabella Rauch (Oregon Health and Science University). *Aim2^−/−^* mice were generated as previously described and housed at the University of Arizona [34].

### 2.3 Cell Culture

Primary bone marrow macrophages (BMMs) were generated by culturing bone marrow isolated from mouse femurs in 10 mLs of DMEM containing 4.5 g/L glucose, L-glutamine, sodium pyruvate (10-013-CV Corning), 20% heat inactivated fetal bovine serum (Gibco), 15% conditioned media from GM-CSF-producing CMG14-12 cells and 1X penicillin-streptomycin (HyClone Thermo) for 7 days. Adherent BMMs were removed from the plates using 0.25% Trypsin-EDTA (Corning) and plated overnight prior to infection.

### 2.4 *In vitro* infection

Primary BMMs were plated at a seeding density of 1.5×10^5^ for ELISA and 3×10^5^ for Western Blot on 48 well and 12 well plates, respectively, in triplicate per genotype. BMMs were infected with *S. anginosus* at a MOI of 10 for 24 hours at 37°C in 200 uL or 1 mL of media without antibiotics for ELISA or Western Blot, respectively. For *S. anginosus* viability studies, *S. anginosus* was heat killed by exposure to 80°C for 30 minutes prior to infecting BMMs. A subset of heat killed *S. anginosus* was plated on blood agar plates and kept at 37°C for 18 hours to ensure no live bacteria remained. WT and *Nlrp3^−/−^* BMMs were treated with 5 μg/mL LPS for 3 hours followed by 7.5 μg/mL of Nigericin (InvivoGen) for 1 hour as a positive control for NLRP3 inflammasome activation. WT and *Aim2^−/−^* BMMs were treated with 5 μg/mL of LPS for 3 hours followed by 1 μg/mL of poly(dA:dT) (InvivoGen) in lipofectamine for 6 hours as a positive control for AIM2 inflammasome activation. For infections also involving pharmacological inhibition of the NLRP3 inflammasome, 10 mM of MCC950 or 0.01% DMSO (vehicle control) was added to each well containing DMEM without FBS for 15 minutes prior to infection. Cells were washed with PBS, supplemented with DMEM containing FBS and infected with *S. anginosus* for 24 hours. After infection or treatment with inflammasome positive control ligands, supernatant or protein was collected and stored at −80°C before assessing cytokine production by ELISA or protein measurements by Western Blot.

### 2.5 *In vivo* Infection

Mice were injected intraperitoneally with 2×10^9^ CFUs of *S. anginosus* suspended in 200 μL of PBS. Animals were weighed at 0, 5, 24, 28 and 48 hours post-infection. After 24 or 48 hours, blood was taken via heart stick and added to a Mini Collect tube (Greiner 450742) for serum isolation per manufacturer’s instructions. Serum was stored at −80°C until used for downstream analysis. Spleen and liver were removed, cut in half and one half was homogenized in 10 μL/mg protein sterile PBS. Serial dilutions of protein homogenates were plated on blood agar plates overnight for CFU counts on the following day. Additional half sections of spleen and liver were quartered, flash frozen in liquid nitrogen and stored at −80°C for protein and RNA assessments.

### 2.6 Crystal Violet Staining and Imaging Quantification

*S. anginosus*-infected BMMs were washed with PBS and fixed with 4% paraformaldehyde for 15 minutes at room temperature. Fixed BMMs were stained with 0.25% crystal violet for 30 minutes at room temperature. Crystal violet was aspirated after the incubation. The cells were washed with water until all stain was removed and air dried for 15 minutes. BMMs were imaged at 20X using ECHO Revolve D2224. BMM areas were quantified using ImageJ software (NIH).

### 2.7 Western Blot

Tissue was homogenized in 10 μL/mg protein of RIPA lysis buffer containing PhosphoSTOP (Roche) and Complete protease inhibitor (Roche). Homogenates were further lysed at 4°C on a tube rotator, then centrifuged at 4°C at 15,000 x g for 10 minutes to pellet the insoluble fraction. Soluble lysate was removed and 4X LDS sample buffer (Life technologies) with 0.5M DL-dithiothreitol (R&D) was added.

Protein was separated on a SDS-polyacrylamide gels (BioRad) and transferred onto nitrocellulose membranes (Cytiva). Membranes were blocked for 1 hour in 5% milk in Tris-Buffered Saline with Tween (TBST) followed by incubation with primary antibodies diluted at 1:1,000 in 5% non-fat milk in TBST overnight at 4°C. Primary antibodies were washed five times with TBST, and membranes were then incubated with secondary antibodies conjugated to horseradish peroxidase diluted at 1:10,000 in 5% non-fat milk in TBST for two hours at room temperature. The membranes were washed with TBST and exposed to SuperSignal West Pico PLUS Chemiluminescent Substrate (ThermoFisher Scientific) for visualization on Blue Devil Film, which was developed using an automatic film processor (OPTIMAX). Protein band intensity was quantified using densitometric analysis via ImageJ software. Primary antibodies used were rabbit mAb anti-COX2 (Cell Signaling 12282S), rabbit mAb anti-NLRC4 (Cell Signaling 12421T), rabbit mAb anti-NLRP3 (Cell Signaling 1510S), rabbit mAb phopsho-p65 (S536) (Cell Signaling 3033T), rabbit mAb anti-p65 (Cell Signaling 8242S), rabbit mAb anti-iNOS (abcam ab283655), rabbit mAb anti-AIM2 E518Z (Cell Signaling 53491S), and anti-β-Actin-HRP (Santa Cruz Biotech A1723). Goat Anti-Rabbit IgG, HRP conjugate (Millipore 12-348) was used as a secondary antibody for all listed primary antibodies with the exception of β-Actin.

### 2.8 ELISA

Enzyme-linked immunosorbent assays (ELISA) were performed on macrophage supernatants and animal serum per manufacturer’s instructions. Briefly, a 96 well ELISA plate was coated with capture antibody overnight at room temperature and washed three times with wash buffer. The plate was blocked with reagent diluent at room temperature for 1 hour and washed three times with wash buffer. Cytokine standards, cell supernatants and animal serum was added to the plate for a 2 hour incubation at room temperature. The plate was washed three times with wash buffer. Detection antibody was added to the plate for a 1 hour incubation at room temperature. The plate was washed three times with wash buffer, and Streptavidin-HRP was added for a 20 minute incubation at room temperature. The plate was washed three times with wash buffer, and substrate solution was added to the plate for a 20 minute incubation at room temperature. Stop solution was added to the plate, and optical density was measured using a microplate reader set to 450 nm. The ELISA kits used included Mouse IL-1 beta/IL-1F (R&D Systems DY401), Mouse TNF-alpha (R&D Systems DY410-05) and Mouse IL-6 (R&D Systems DY406-05).

### 2.9 Bacteria Killing Assay

Primary BMMs were plated in triplicate on 12 wells plates at a seeding density of 2×10^5^ cells/well and allowed to adhere overnight at 37°C. BMMs were infected with *S. anginosus* at a MOI of 10 for 30 minutes. BMMs were then washed with PBS to remove external bacteria and treated with 150 ug/mL of gentamycin for 30 minutes to neutralize any non-internalized bacteria before a final PBS wash. The BMMs were then incubated in antibiotic-free media for 2.5 or 24 hours. The BMMs were given a final PBS wash and lysed in 100 uL of sterile water with scraping. The lysates were serially diluted five times, plated on blood agar plates and incubated at 37°C overnight for CFU counts the following day. A subset of macrophages were washed and lysed after the initial 30 min bacterial infection to determine the total number of internalized bacteria.

### 2.11 LDH Assay

*S. anginosus*-induced macrophage cell death was assessed using a Lactate dehydrogenase (LDH) assay according to manufacturer’s protocol (AbCam ab65393). In brief, primary BMMs were plated in triplicate on 48 well plates at a seeding density of 1.5×10^5^ cells/well and incubated overnight at 37°C. BMMs were infected with *S. anginosus* at a MOI of 1, 10, 25, 50, 75 and 100 for 24 hours, and supernatant was taken. A subset of uninfected BMMs were incubated with lysis buffer as a positive control. A total of 10 μl of supernatant was mixed with 100 μl of LDH reaction mix and incubated at room temperature for 30 mins. The samples were then read at OD 450 nm using a microplate reader. The %LDH release was calculated as the sample OD 450 nm divided by the positive control OD 450 nm.

### 2.12 Statistical Analysis

The data were expressed as the mean +/− the standard error of the mean (SEM) of values obtained from experiments. Statistical analysis was performed using GraphPad Prism Software using ANOVA or unpaired t test. P values of <.05 were considered statistically significant.

## 3. Results

### 3.1. *S. anginosus* infection elicits an inflammatory response in bone marrow macrophages *in vitro*

Our previous work indicates that *S. anginosus* induces a robust proinflammatory response accompanied by distinct morphologic changes in RAW 264.7 cells after 6 hour infection [32]. In order to assess the inflammatory potential of *S. anginosus* in more physiologically relevant primary cells, we generated bone marrow-derived macrophages (BMMs) from C57BL/6 (wild type) mice and infected them with *S. anginosus* at a MOI of 10 for 6 and 24 hours. After 6 hours of *S. anginosus* infection, primary BMMs displayed an elongated phenotype with a significantly smaller area than uninfected cells (**Fig 1a**), similar to *S. anginosus*-infected RAW 264.7 cells [32]. However, following 24 hours post-infection, BMMs displayed an enlarged phenotype and lost their elongated phenotype (**Fig 1a**). These findings suggest that primary BMMs display morphological changes reminiscent of an activated state in response to *S. anginosus* infection. Therefore, we next examined proinflammatory signaling and downstream proteins in *S. anginosus*-infected BMMs. This included phosphorylated levels of the NF-κB family member p65 and expression of iNOS and COX2, which are responsible for production of proinflammatory effectors nitric oxide and prostaglandins, respectively. After 6 hours of infection with *S. anginosus*, BMMs displayed a significant increase phospho-p65, iNOS and COX2 (**Fig 1b**). These pro-inflammatory markers remained elevated at 24 hours post infection (**Fig 1c**). The NF-κB family of transcription factors are required for the production of proinflammatory cytokines. Thus, we infected BMMs with *S. anginosus* for 24 hours and measured the levels of proinflammatory cytokines TNF⍰, IL-1β and IL-6 in supernatants. TNF⍰, IL-1β and IL-6 were significantly upregulated after 24 hour *S. anginosus* infection (**Fig 1d**). These results indicate that *S. anginosus* induces a robust pro-inflammatory response in primary BMMs *in vitro*.

**Figure 1.**
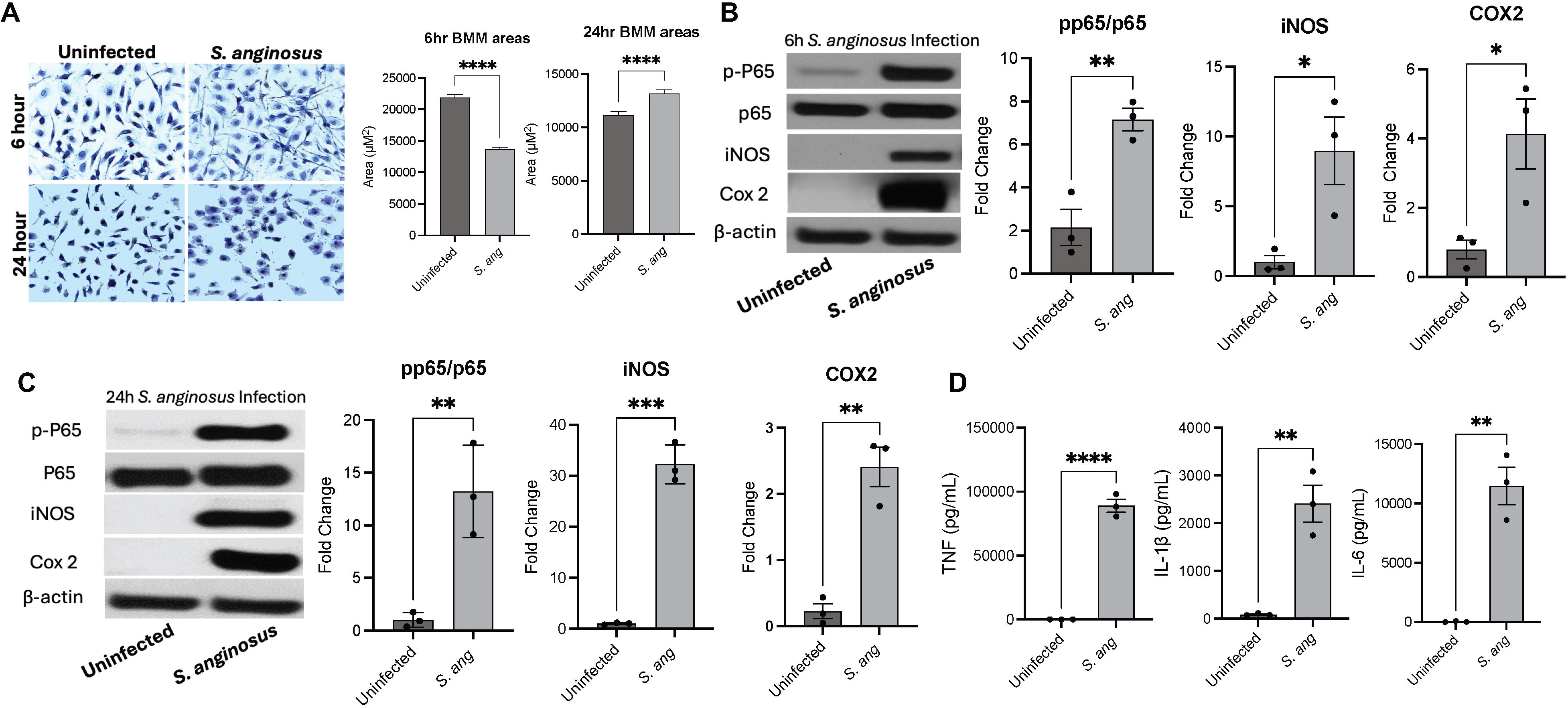
*S. anginosus* infection elicits an inflammatory response in bone marrow macrophages *in vitro*: a) 40x images of BMMs infected with *S. anginosus* for 6h and 24h at a MOI of 10 and uninfected, stained with crystal violet. Quantification of 2000-3000 cell areas per condition. b) western blot and densitometry of phospho-p65, p65, iNOS and COX2 from BMMs uninfected and infected for 6h at a MOI of 10. Quantification of three biological replicates. c) western blot and densitometry of phospho-p65, p65, iNOS and COX2 from BMMs uninfected and infected for 24h at a MOI of 10. Quantification of three biological replicates. d) ELISA of TNFα, IL-1β and IL-6 from BMM supernatant uninfected and infected for 24h at a MOI of 10. Quantification of three biological replicates. Data is presented as mean +/− SEM. Stats quantified using an unpaired T test through GraphPad. *P<.05, **P<0.001, ***P<0.0005, ****P <0.0001 in comparison to uninfected conditions.

### 3.2. *S. anginosus* viability is required to induce fulminate inflammatory responses in bone marrow macrophages

We next determined the viability requirements of *S. anginosus* to induce a proinflammatory response in BMMs. To test this, we inactivated *S. anginosus* by heat exposure at 80°C for 30 minutes in order to maintain the structural integrity of *S. anginosus*, which should retain the capacity to activate surface PRRs on BMMs without releasing additional factors resulting from bacterial lysis. To confirm inactivation, we plated a subset of heat inactivated *S. anginosus* on blood agar plates, which resulted in no measurable CFUs. We then exposed BMMs to either live or heat-killed (HK) *S. anginosus* for 24 hours and assessed BMM morphologic changes after crystal violet staining by microscopy. Both live and HK *S. anginosus* induced an enlarged phenotype in BMMs, however HK *S. anginosus* induced BMM enlargement at a significantly reduced capacity compared to live *S. anginosus* (**Fig 2a**). Moreover, live *S. anginosus* infection induced a greater elongated phenotype in BMMs compared to HK *S. anginosus* infection. These findings indicate that *S. anginosus* viability may be partially responsible for its ability to activate macrophages. Next, to determine if *S. anginosus* viability is required for its ability to induce proinflammatory pathways in macrophages, we assessed levels of phosphorylated p65, iNOS and COX2 in BMMs infected with live or HK *S. anginosus*. While HK *S. anginosus* induced similar levels of phopho-p65 as live *S. anginosus*, HK *S. anginosus* displayed a reduced capacity to induce iNOS and COX2 in BMM*s* (**Fig 2b**). Additionally, BMMs infected with HK *S. anginosus* produced significantly less proinflammatory cytokines such as IL-1β and TNF⍰ compared to BMMs given live *S. anginosus* (**Fig 2c**). These findings indicate that *S. anginosus* exposure leads to activation of the NF-κB pathway independent of bacterial viability. This is likely mediated by TLRs that recognize microbial products that are present on the surface of *S. anginosus,* irrespective if live or heat killed. However, infection with live *S. anginosus* was needed for a strong proinflammatory response and induction of important downstream proinflammatory proteins and cytokines. Overall, this indicates that live *S. anginosus* is needed for the complete inflammatory response initiated by primary BMMs and the induction of proinflammatory proteins and cytokines.

**Figure 2.**
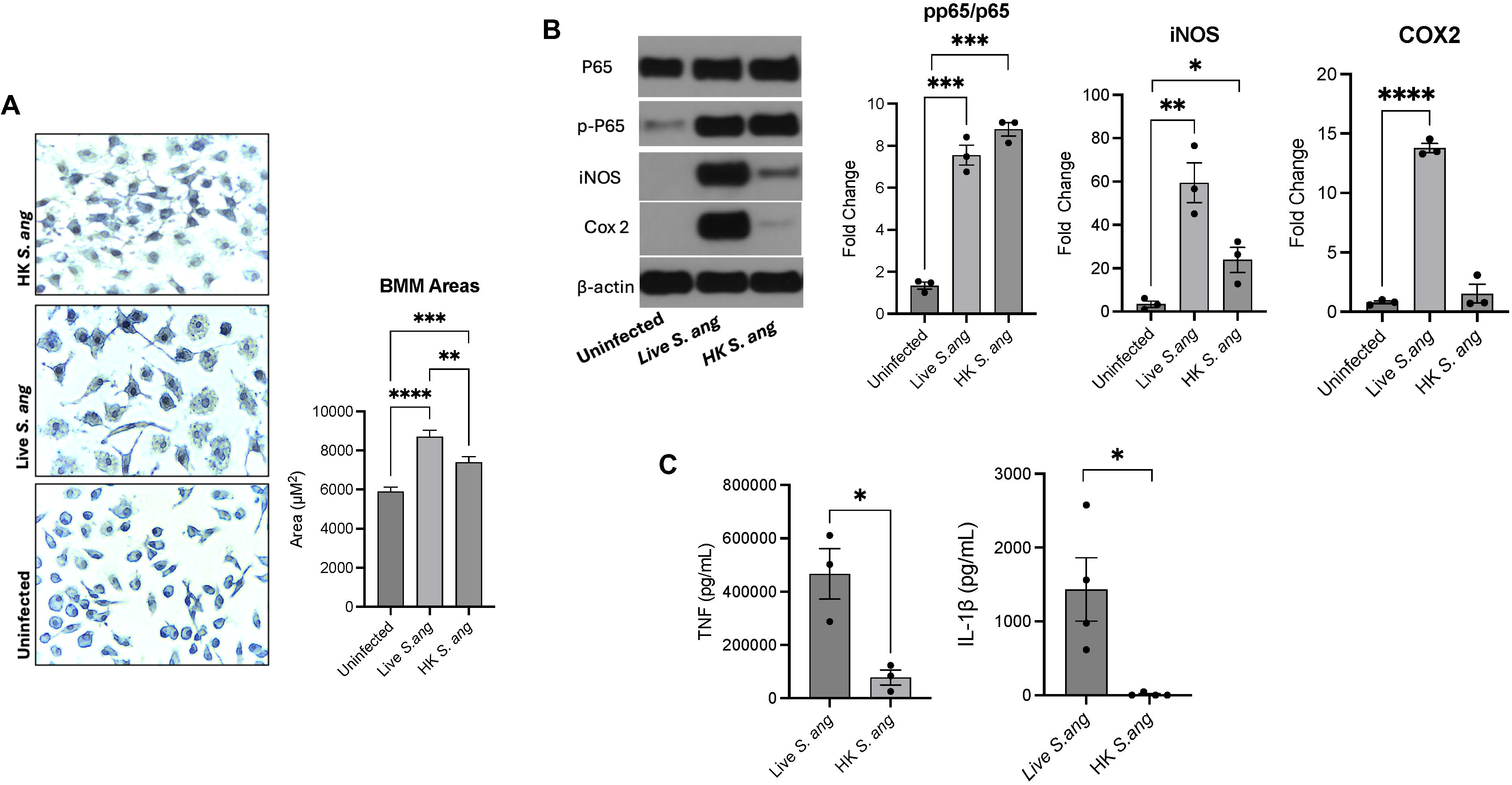
*S. anginosus* viability is required to induce fulminate inflammatory response in bone marrow macrophages: a) 40x images of BMMs uninfected or infected for 24h with live or heat killed *S. anginosus* at a MOI of 10, stained with crystal violet. Quantification of 4000-6000 cell areas per condition. b) western blot and densitometry of phospho-p65, p65, iNOS and COX2 from BMMs uninfected or infected for 24h with live or heat killed *S. anginosus* at a MOI of 10. Quantification of three biological replicates. c) ELISA of TNFα and IL-1β from BMM supernatant uninfected or infected for 24h with live or heat killed *S. anginosus*. Quantification of three biological replicates. Data is presented as mean +/− SEM. Stats quantified using an unpaired T test through GraphPad. *P<.05, **P<0.001, ***P<0.0005 ****P <0.0001 in comparison to uninfected conditions.

### 3.3. *S. anginosus* replicates within infected macrophages and promotes macrophages cell death at high MOI

*S. anginosus* viability was required to promote production of downstream proinflammatory proteins and cytokines from macrophages. This lead us to postulate that live *S. anginosus* presents an insult to infected macrophages that necessitate this robust inflammatory response. For example, while bacteria are typically neutralized following phagocytosis, other oral streptococcus species including *S. gordonii* replicate intracellularly, which triggers production of the potent inflammatory cytokine IL-1β [26]. In order to better understand how macrophages respond to *S. anginosus*, we next assessed the ability of *S. anginosus* to persist within BMMs using a bacterial killing assay as done previously with *S. gordonii* [26]. Accordingly, we infected BMMs with *S. anginosus* for 30 min, then treated BMMs with gentamicin to neutralize any non-internalized bacteria and incubated the BMMs for an additional 2.5 or 24 hrs. BMMs were lysed, and lysates were then plated on blood agar plates to determine CFU counts. A subset of BMMs were lysed after the initial 30 min infection to determine the total number of internalized bacteria. We observed an initial decrease in internalized *S. anginosus* at 2.5 hours post-infection relative to the 30 min baseline measurements (representing the total input of bacterial). However, at 24 hrs post-infection, *S. anginosus* CFUs were significantly increased compared to the initial 30 min time points (**Fig 3b**). These results suggest that macrophages fail to neutralize phagocytosed *S. anginosus,* resulting in replication of *S. anginosus* in infected BMMs. We next asked if *S. anginosus* in turn, promotes macrophage cell death during infection. Our prior work in RAW264.7 cells suggests *S. anginosus* does not induce any measurable cell death at an MOI 10, [32], however this has not been confirmed in primary macrophages. To determine if *S. anginosus* infection induces cell death in primary macrophages, we measured LDH release from BMMs infected at range of MOIs from 1-100. BMMs infected at MOIs of 1 and 10, the maximum MOI used in this study, had LDH release similar to that of uninfected cells, indicating *S. anginosus* induced a robust inflammatory response without causing measurable cell death in macrophages. However, exposure of BMMs with *S. anginosus* at MOIs of 25 and higher lead to a significant release of LDH, indicating *S. anginosus* was capable of inducing cell death, but only at high MOIs (**Fig 3c**). Because our previous experiments used infections at an MOI of 10 to assess *S. anginosus*-induced inflammatory responses in BMMs, we next determined if *S. anginosus*-induced production of proinflammatory cytokines (i.e., IL-1β) was responsive to increasing MOIs. However, BMMs infected with *S. anginosus* at MOIs of 10, 25 and 50 secreted a similar amounts of IL-1β (**Fig 3d**). Together, this suggests that macrophages fail to effectively neutralize *S. anginosus* following phagocytosis, and that *S. anginosus* is instead capable of replicating within BMMs. Moreover, despite replicating within macrophages, *S. anginosus* does so without promoting macrophage cell death unless at a high (>10) MOI.

**Figure 3.**
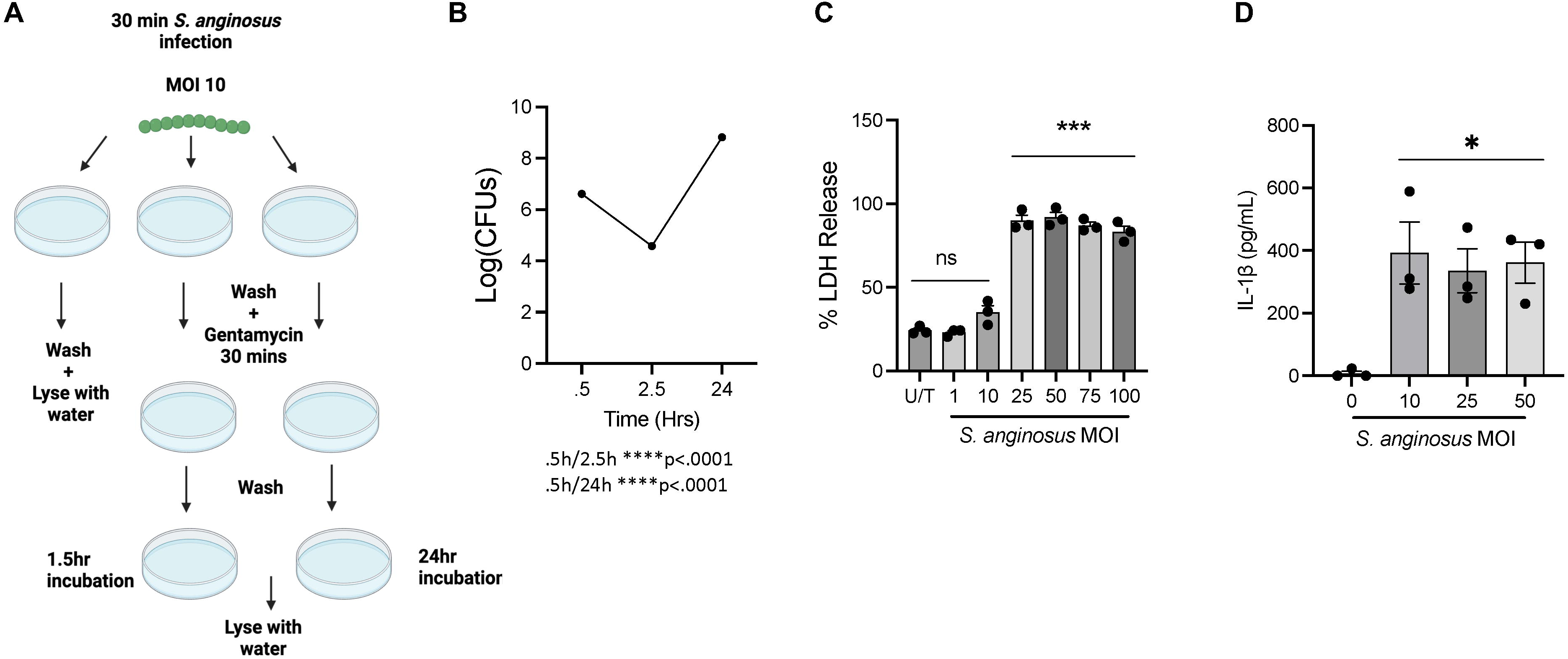
*S. anginosus* replicates within macrophages and promotes macrophages cell death at high MOI: a) schematic of the bacteria killing assay b) bacteria killing assay looking at internalized *S. anginosus* after 30 min, 2.5h and 24h infection of BMMs. Shown as an average of three biological replicates per time point. c) LDH assay of BMMs uninfected and infected at a MOI of 1, 10, 25, 50, 75 and 100 with *S. anginosus*. Percent LDH released quantified as experimental value divided by the positive control. Quantification of three biological replicates. d) ELISA of IL-1β from BMM supernatant uninfected or infected at a MOI of 10, 25 and 50 for 24 hours. Quantification of three biological replicates. Data is presented as mean +/− SEM. Stats quantified using an unpaired T test through GraphPad. *P<.05, ***P<0.0005, ****P <0.0001 in comparison to uninfected conditions.

### 3.4. *S. anginosus* activates the NLRP3 inflammasome

IL-1β is a potent proinflammatory cytokine that is often produced by macrophages in response to intracellular infection [35]. Production of IL-1β is mediated by a 2-step process involving: 1) a priming step whereby NF-κB activation leads to transcription and production of inactive pro-IL-1β; and 2) activation and oligomerization of the multi-protein complex termed the inflammasome, which activates IL-1β and promotes its release from the cell [36]. Several inflammasome sensors are triggered by other streptococcus species, including NLRP3, NLRC4 and AIM2 [24], [31], [28]. Indeed, the protein expression of NLRP3, NLRC4 and AIM2 were all upregulated in response to *S. anginosus* infection (**Fig 4a**), indicating *S. anginosus* induces the expression of multiple inflammasome sensors. To determine if *S. anginosus*-induced IL-1β production was inflammasome dependent, we infected BMMs lacking the common inflammasome adapter protein ASC (*Asc^−/−^*) with *S. anginosus* for 24 hours and assessed IL-1β. *S. anginosus*-infected *Asc^−/−^* BMMs produced normal levels of TNF⍰, indicating that NF-κB activation was intact in these BMMs (**Fig. 4b**). However, there was significant diminishment of IL-1β production in *Asc^−/−^* BMMs, indicating that *S. anginosus-*induced IL-1β production is inflammasome dependent (**Fig 4b**). To determine which inflammasome sensor(s) are triggered by *S. anginosus* to induce IL-1β production, we infected BMMs lacking NLRP3, NLRC4, or AIM2. *Nlrp3^−/−^*, *Nlrc4^−/−^* and *Aim2^−/−^*BMMs still produced TNF⍰ (**Fig 4c, d, e**). In addition, while *Nlrc4^−/−^* and *Aim2^−/−^* BMMs produced similar levels of IL-1β compared to wild type BMMs, *Nlrp3^−/−^* BMMs had significantly diminished, but not complete loss of IL-1β production in comparison to wild type BMMs (**Fig 4c, d, e**). To confirm that *S. anginosus* was acting through the NLRP3 inflammasome using a pharmacological approach, we infected wild type BMMs in the presence of DMSO vehicle or 10 mM of the NLRP3 inhibitor, MCC950, for 24 hours. IL-1β was significantly diminished in wild type BMMs treated with MCC950 in comparison to those treated with the vehicle control (**Fig 4f**). This implicates that *S. anginosus* activates the inflammasome in part, through triggering NLRP3 to induce IL-1β *in vitro*.

**Figure 4.**
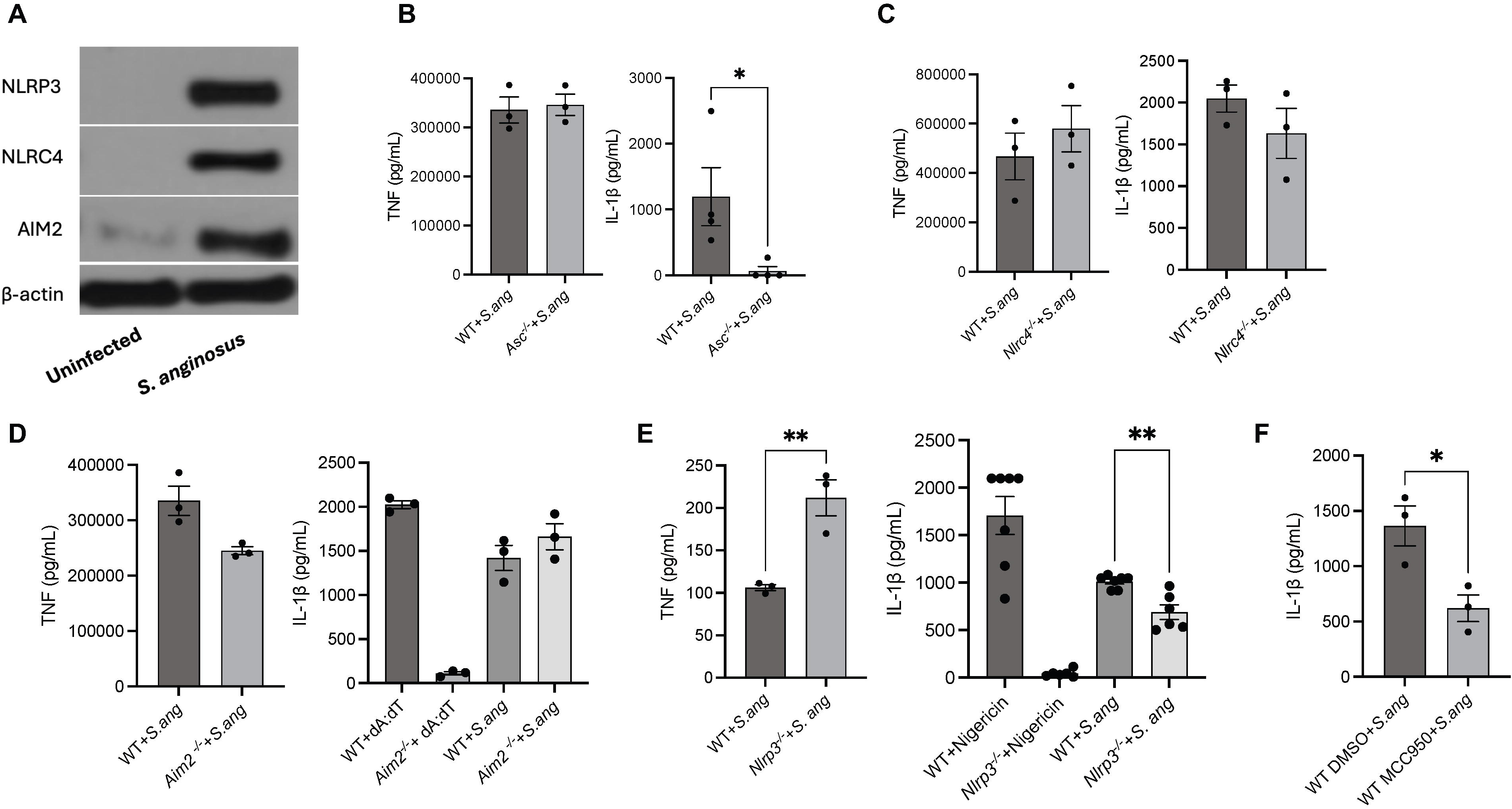
*S. anginosus* activates the NLRP3 inflammasome: a) western blot of NLRP3, NLRC4 and AIM2 from BMMs uninfected or infected for 24h with *S. anginosus* at a MOI of 10. b) ELISA of TNFα and IL-1β from WT and *Asc^−/−^* BMM supernatant uninfected or infected for 24h. Quantification of three biological replicates. c) ELISA of TNFα and IL-1β from WT and *Nlrc4^−/−^* BMM supernatant uninfected or infected for 24h with *S. anginosus*. Quantification of three biological replicates. d) ELISA of TNFα and IL-1β from WT and *Aim2^−/−^* BMM supernatant infected or uninfected for 24h. 1h LPS prime followed by 6h treatment of poly dA:dT was used as a positive control. Quantification of three biological replicates. e) ELISA of TNFα and IL-1β from WT and *Nlrp3^−/−^* BMM supernatant infected or uninfected for 24h. 1h LPS prime followed by 3h nigericin treatment was used as a positive control. Quantification of three biological replicates. f) ELISA of IL-1β from WT BMM supernatant uninfected or infected for 24h with or without 10mM NLRP3 inhibitor, MCC950. Data is presented as mean +/− SEM. Stats quantified using an unpaired T test through GraphPad. *P<.05, **P<0.001 in comparison to uninfected conditions. Data without stats are ns.

### 3.5. *S. anginosus* elicits an acute inflammatory response *in vivo*

To determine if *S. anginosus* induces a proinflammatory response *in vivo*, we infected wild type mice with 2×10^9^ CFUs of *S. anginosus* via intraperitoneal injection (i.p.), an approach used to assess the inflammatory capacity of similar streptococcus species in vivo [37]. After 24 hours of infection, wild type mice displayed significant weight loss, which largely recovered by 48 hours, but remained significantly lower than the initial starting weights. In addition, *S. anginosus* was detectable in the spleen and liver of wild type mice 24 hours-post infection, indicating bacterial dissemination occurred to these organs (**Fig. 5b**). To determine the role of the inflammasome during the host response to *S. anginosus* infection *in vivo*, we next infected *Asc^−/−^* mice with 2×10^9^ CFUs of *S. anginosus*. *Asc^−/−^* mice displayed significantly reduced weight loss at 24 hours compared to similarly-infected wild type mice, which was almost completely resolved by 48 hrs (**Fig 5a**). Compared to wild type mice, *Asc^−/−^* mice displayed similar dissemination of *S. anginosus* to the spleen and slightly higher liver dissemination, however this level did not reach significance (**Fig 5b**). We next assessed the level of inflammation in disseminated tissue by measuring phospho-p65 expression in spleen and liver homogenates. Expression of phopho-p65 was similar in the spleen of *S. anginosus*-infected wild type and *Asc^−/−^* mice after 24 hours of infection (**Fig 5c, Supplemental Fig 1A**). Additionally, we observed similar levels of IL-1β in the serum of both wild type and *Asc^−/−^* mice at 24 hours post infection (**Fig 5d**), suggesting inflammasome activation is dispensable for these systemic levels of IL-1β. By 48 hours post infection, *S. anginosus* was not detectable in the spleen or liver of wild type and *Asc^−/−^* mice, indicating that wild type mice resolved this infection, which was independent of the inflammasome (data not shown). Similar to 24 hour values, we detected comparable levels of phospho-p65 in the spleen of wild type and *Asc^−/−^* mice at 48 hours (**Fig 5e, Supplemental Fig 1B**). However, there was a significant increase in phospho-p65 in the livers of *Asc^−/−^* mice in comparison to wild type animals (**Fig 5e, Supplemental Fig 1C**), however no significant differences in serum IL-1β was detected in *Asc^−/−^* mice in compared to wild type mice following 48 hours of *S. anginosus* infection (**Fig 5f**). These findings indicate that *S. anginosus* induces an acute pro-inflammatory response *in vivo* that is resolved by 48 hours and is partially driven by the inflammasome.

**Figure 5.**
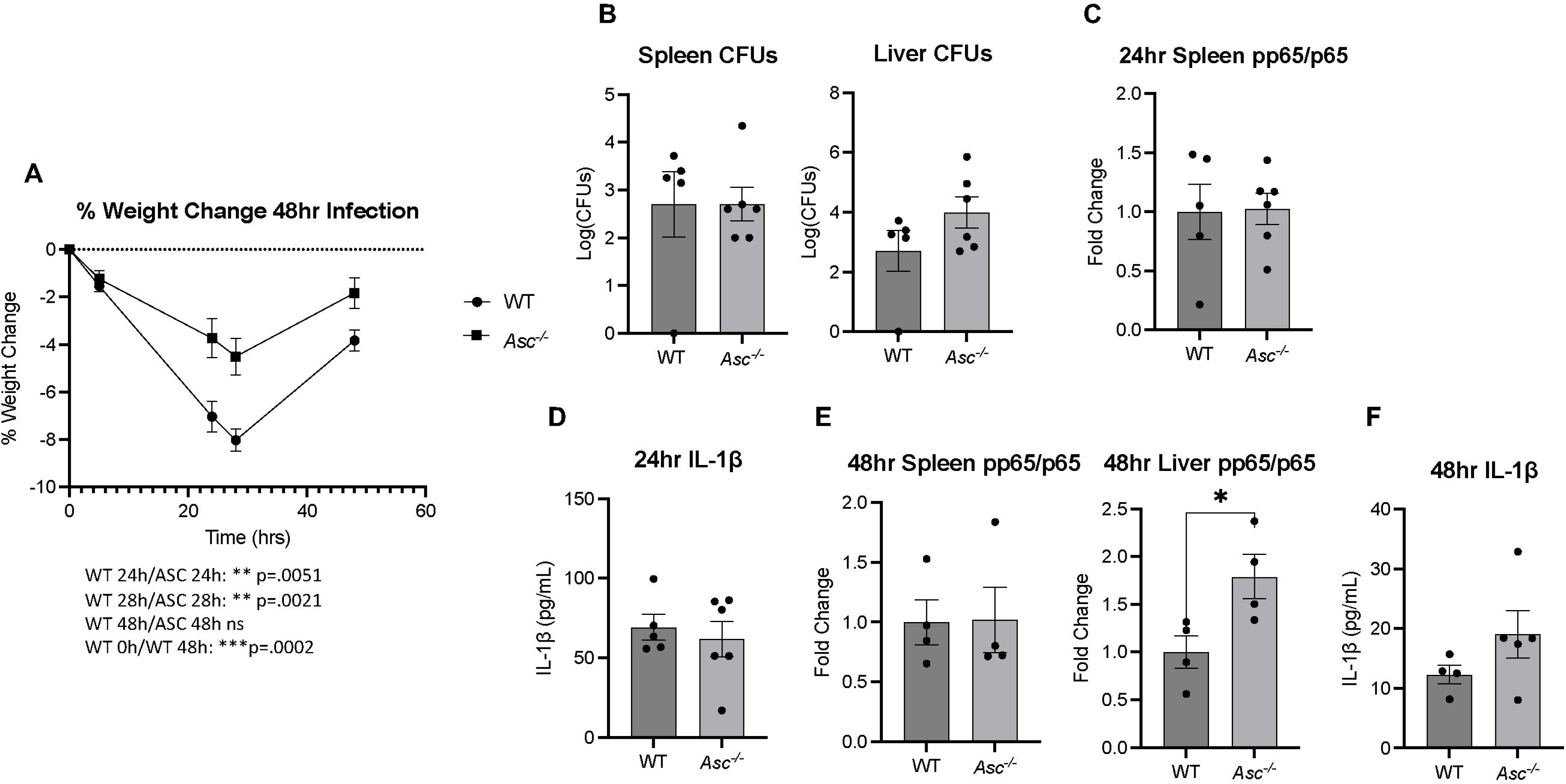
*S. anginosus* elicits an inflammatory response *in vivo*: a) percent weight change of WT or *Asc^−/−^* mice infected by IP injection with 2×10^9^ CFUs of *S. anginosus* for 48 hours. WT n=8, *Asc^−/−^* n=9. b) Log(CFUs) of *S. anginosus* from the spleen and liver of WT and *Asc^−/−^* mice 24h post infection. c) densitometry of phospho-p65/total p65 from the spleen of WT and *Asc^−/−^*mice 24h post infection. d) ELISA of serum IL-1β from WT and *Asc^−/−^* mice 24h post infection. e) densitometry of phospho-p65/total p65 from the spleen and liver of WT and *Asc^−/−^* mice 48h post infection. f) ELISA of serum IL-1β from WT and *Asc^−/−^* mice 48 post infection. Data is presented as mean +/− SEM. Stats quantified using an unpaired t test through GraphPad. *P<.05 in comparison to uninfected conditions. Data without stats are ns.

### 3.6. NLRP3 promotes an Inflammatory response to *S. anginosus in vivo*

Our results indicate that compared to wild type mice, *Asc^−/−^*mice display reduced morbidity in response to *S. anginosus* infection *in vivo*, indicating that the inflammasome plays a partial role during the inflammatory response to *S. anginosus in vivo* (**Fig. 5**). Moreover, NLRP3 was found to be an important inflammasome sensor that responds to *S. anginosus* infection *in vitro* (**Fig. 4**). To verify if the NLRP3 inflammasome sensor is responsible for mediating an inflammatory responses to *S. anginosus in vivo*, we infected wild type and *Nlrp3^−/−^* mice with 2×10^9^ CFUs *S. anginosus* via i.p. injection. We also included *Aim2^−/−^* mice as an additional inflammasome control. At 24 hours post infection, *Nlrp3^−/−^*mice, displayed reduced weight loss in comparison to wildtype mice (**Fig 6a**), similar to our observations with *Asc^−/−^* animals. Conversely, *S. anginosus*-infected *Aim2^−/−^*mice displayed greater weight loss than wild type mice (**Fig 6a**), suggesting AIM2 may play an inflammasome-distinctive role during *S. anginosus* infection *in vivo.* However, we observed no significant differences in spleen or liver dissemination in *Nlrp3^−/−^* and *Aim2^−/−^* mice in comparison to wild type animals (**Fig 6b**). Consistent with these findings, no differences in phospho-p65 were detected in the spleens and livers of *S. anginosus*-infected *Nlrp3^−/−^* and *Aim2^−/−^* mice in comparison to wild type animals (**Fig 6c, Supplemental Fig 2A+B**). Serum IL-1β was also similar in infected wild type, *Nlrp3^−/−^* and *Aim2^−/−^* mice (**Fig 6d**). Similar to *Asc^−/−^* and wild type mice, we failed to detect dissemination of *S. anginosus* to the spleen and liver of *Aim2^−/−^*and *Nlrp3^−/−^* mice 48 hrs post infection (data not shown). However, at 48 hours post-infection, a significant increase in phospho-p65 was detected in the spleen of *Aim2^−/−^* mice in comparison to similarly-infected wild type mice, with no significant differences detected in *Nlrp3^−/−^* mice (**Fig 6e, Supplemental Fig 2C**). Serum IL-1β levels were similarly diminished in wild type, *Aim2^−/−^*and *Nlrp3^−/−^* mice after 48 hrs infection compared to levels observed at 24 hours (**Fig 6f**), further indicating resolution of infection occurred independent of the inflammasome *in vivo.* These findings indicate that the NLRP3 plays a partial role in promoting the morbidity associated with *S. anginosus* infection *in vivo*.

**Figure 6.**
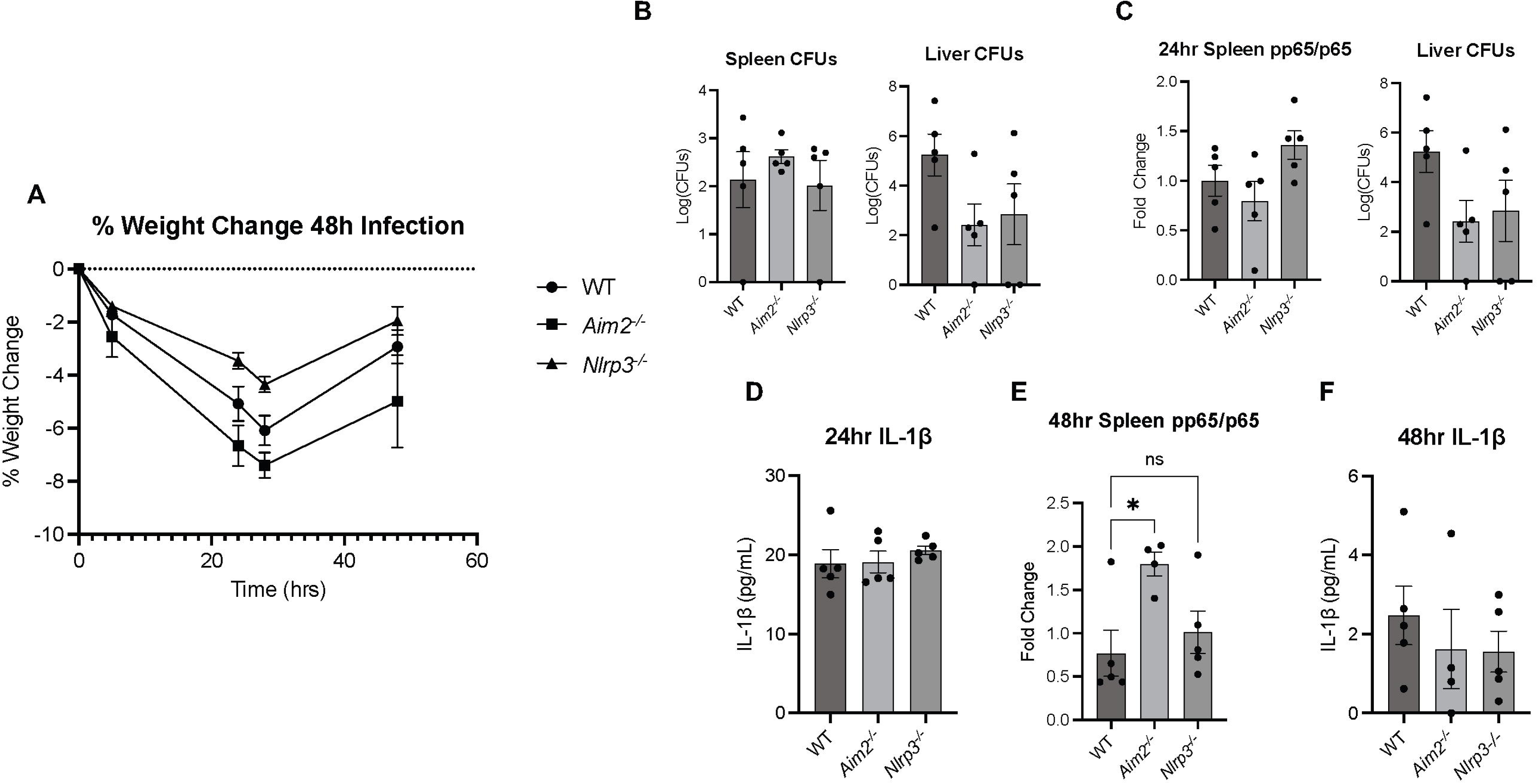
NLRP3 promotes an Inflammatory response to *S. anginosus in vivo*: a) percent weight change of WT, *Aim2^−/−^*or *Nlrp3^−/−^* mice infected by IP injection with 2×10^9^ CFUs *S. anginosus* for 48 hours. WT n=5, *Aim2^−/−^* n=4, *Nlrp3^−/−^* n=5. b) Log(CFUs) of *S. anginosus* in the spleen and liver of WT, *Aim2^−/−^* and *Nlrp3^−/−^* mice 24h post infection. c) densitometry of phospho-p65/total p65 from the spleen and liver of WT, *Aim2^−/−^* and *Nlrp3^−/−^* mice 24h post infection. d) ELISA of serum IL-1β from WT, *Aim2^−/−^* and *Nlrp3^−/−^* mice 24h post infection. e) densitometry of phospho-p65/total p65 from the spleen of WT, *Aim2^−/−^* and *Nlrp3^−/−^*mice 48h post infection. f) ELISA of serum IL-1β from WT, *Aim2^−/−^* and *Nlrp3^−/−^* mice 48h post infection. Data is presented as mean +/− SEM. Stats quantified using an unpaired t test through GraphPad. *P<.05 in comparison to uninfected conditions. Data without stats are ns.

## 4. Discussion

Although *S. anginosus* is a common commensal of the oral cavity, it’s abundance is reportedly increased during OSCC and it can directly impact the epithelium to drive tumor formation and progression [38], [39]. *S. anginosus* also promotes gastric cancer progression, which is associated with elevated inflammatory cell infiltration and the induction of the chemokines CCL20 and CXCR2 *in vivo* [16], yet how *S. anginosus* interacts with the innate immune system to drive the inflammatory responses during tumor initiation and progression remains unclear. Previously, we found that *S. anginosus* induces the inflammatory cytokines IL-1β, TNF⍰ and IL-6 in the RAW 264.7 macrophage cell line [32]. Additionally, we found that *S*. *anginosus* induces the expression of the cytosolic sensor NLRP3 and causes distinct morphological changes in RAW264.7 cells reminiscent of cellular activation [32]. While our previous work established a cellular link between *S. anginosus* and innate immune activation, the molecular mechanism behind this relationship remained to be investigated. In this study, we expanded on our previous findings using primary BMMs for greater physiological relevance and first confirmed that *S. anginosus* promotes a robust pro-inflammatory response in primary macrophages. This includes elevated NF-κB signaling, increased proinflammatory cytokine production (i.e., TNFα, IL-6 and IL-1β) and increased protein expression of inflammatory mediators COX2 and iNOS in *S. anginosus*-infected BMMs. Additionally, we observed that macrophages exposed to *S. anginosus* displayed an increase in size after 24 hour infection, consistent with cellular activation and consistent with our previous findings in RAW264.7 cells [32]. We next focused our attention on understanding the mechanism by which *S. anginosus* drives macrophage activation, starting with the question: is bacterial viability required for the observed pro-inflammatory responses? We found that *S. anginosus* viability was dispensable for the early events of inflammatory signaling as heat-killed *S. anginosus* induced macrophage p65 activation equally as well as live *S. anginosus* following 24 hrs of infection. However, *S. anginosus* viability was essential for the induction of several downstream pro-inflammatory mediators like iNOS and COX2. Additionally, heat-killed *S. anginosus* failed to induce the production of the inflammasome-associated cytokine IL-1β. These findings lead us to explore the possibility that macrophages may fail to effectively neutralize live *S. anginosus* following phagocytosis, which could result in an enhanced proinflammatory response compared to inactivated *S. anginosus*. Indeed, when we infected BMMs with *S. anginosus* and assessed the CFUs of internalized bacteria 24 hours later, we recovered significantly greater CFUs than the initial amount that was used to infect the BMMs. This finding indicated that *S. anginosus* persisted within macrophages. However, it is unclear if *S. anginosus* completely evades macrophage-mediated killing or if the rate of *S. anginosus* replication within macrophages outpaced the rate of bacterial killing. Nonetheless, these findings indicated that macrophages fail to effectively neutralize *S. anginosus.* The initial inflammatory signaling (i.e., NF-κB) induced by the inactivated *S. anginosus* likely occurred through activation of surface PRRs (e.g., TLR2), which recognize structural components on a wide range of microbes irrespective of their replication potential. However, many traditional pathogens often drive more robust and/or persistent inflammatory response through a variety of actions including intracellular replication. Indeed, several other streptococcus species, including *S. gordonii*, *S. aureus* and *S. typhimurium,* can survive within macrophages [26] [40], [41]. While the mechanism of infiltration and survival within macrophages of these other streptococcus species types have been elucidated, how *S. anginosus* accomplishes this remains to be investigated

The innate immune system often relies upon intracellular PPRs, including the inflammasome-forming NOD-like receptor (NLR) proteins, to recognize and mount appropriate responses to intracellular pathogens. The inflammasome is a multi-protein complex consisting of a sensor (e.g., NLRP3 and AIM2), the adapter protein ASC and the enzyme Caspase-1 that, upon activation by a variety of microbial- and damage-associated ligands, leads to IL-1β production and potentially pyroptosis [17]. Because *S. anginosus* persisted within BMMs and induced the inflammasome-dependent cytokine IL-1β, we decided to investigate the requirement of the inflammasome during *S. anginosus* recognition and if *S. anginosus* induces pyroptosis. Indeed, we found that *S. anginosus* failed to induce IL-1β production from macrophages lacking the common inflammasome adapter protein ASC, confirming the involvement of the inflammasome during this process. Additionally, we saw that macrophages infected with *S. anginosus* at the MOI used throughout this study (i.e., 10) released similar levels of LDH as uninfected BMMs, indicating that *S. anginosus* infection at this MOI doesn’t lead to significant cell death. Interestingly, MOIs at 25 and greater caused significant LDH release, while IL-1β secretion remained similar across all MOIs. This indicates that lower levels of *S. anginosus* are sufficient to maintain proinflammatory cytokine production from BMMs without inducing cell death, which in turn may prolong inflammation and exacerbate disease. Additionally, this finding may indicate that *S. anginosus* does not cause a substantial release of damage-associated ligands as other cell-death-inducing pathogenic species of bacteria, which are detected by intracellular PRRs.

Because IL-1β production from *S. anginosus* infection was inflammasome dependent, we investigated if *S. anginosus* was detected by one or multiple inflammasome sensors linked to the recognition of other streptococcus species including NLRP3, AIM2 and NLRC4. NLRP3 can recognize and be activated by a wide variety of PAMPs and DAMPs including extracellular ATP, bacterial toxins like nigericin, reactive oxygen species and changes in ion influx [42]. AIM2 recognizes cytosolic dsDNA from microbes and mammalian cells [43]. NLRC4 traditionally recognizes flagellum, although it has been linked to the recognition of some bacteria lacking flagella, as well as other DAMPs and PAMPs such as lysophosphatidylcholine and retrotransposon RNAs [31] [29] [44]. Here we found BMMs missing NLRP3 had a significant reduction in IL-1β secretion. Additionally, pharmacological inhibition of NLRP3 in wild type BMMs using MCC950 resulted in a similar reduction in IL-1β secretion during *S. anginosus* infection. This is not surprising as NLRP3 is responsible for recognizing infection of many streptococcus species, including *S. pneumoniae*, *S. suis* and *S. pluranimalium* [45] [46], [47]. Overall, these findings implicate the NLRP3 inflammasome during the recognition of *S. anginosus.* Although IL-1β was significantly reduced in *S. anginosus*-infected NLRP3 deficient BMMs, these levels were not as drastic as what was observed in ASC deficient BMMs. Therefore, we cannot rule out the involvement of other and/or multiple inflammasome sensors. For example, NLRP6 recognizes *S. pneumoniae* [25], and multiple inflammasomes (AIM2, NLRP3 and NLRC4) are activated by *S. mutans* in human THP-1 macrophages [48]. However, we can conclude that AIM2 and NLRC4 do not contribute to *S. anginosus*-driven IL-1β since AIM2- and NLRC4-deficient macrophages presented similar levels of IL-1β as their wild type counterparts. As *S. anginosus* has no known flagella, it is not entirely surprising that the NLRC4 inflammasome was not involved in this process. Moreover, NLRC4 can also form an inflammasome independently of ASC, yet ASC was required for *S. anginosus* induced-IL-1β.

However, it was somewhat surprising that AIM2 was dispensable for *S. anginosus*-induced inflammasome activation. This suggests that *S. anginosus* DNA likely does not access the cytosol of infected BMMs during the first 24 hours of infection and therefore may indicate that *S. anginosus* replicates in endosomal compartments while evading killing by macrophages. Additionally, our results suggest that *S. anginosus* likely triggers inflammasome activation through the generation of NLRP3-activating DAMPS and/or through virulence factors. Future studies will focus on characterizing the localization of *S. anginosus* within intracellular compartments of BMMs and identifying how *S. anginosus* triggers NLRP3.

After establishing a mechanistic role for NLRP3 inflammasome during *S. anginosus* infection *in vitro*, we next explored the inflammatory response to *S. anginosus* infection *in vivo*. First, we infected wild type mice with *S. anginosus* and tracked disease progression over 48 hours. We observed significant weight loss in animals at 24 hours post-infection, which was accompanied by bacterial dissemination to the spleen and liver. By 48 hours post-infection, the animals largely recovered but still displayed significant weight loss compared to the initial values observed prior to infection. Moreover, we failed to detect any measurable *S. anginosus* levels in liver and spleen at the 48 hour time point. These findings indicate that *S. anginosus* induces an acute, but self-limiting proinflammatory response during an intraperitoneal infection model. To determine if the inflammasome plays a role during *S. anginosus* infection *in vivo*, we repeated these infections using *Asc^−/−^* mice. Compared to wild type controls, *Asc^−/−^* mice displayed significantly less weight loss at the 24 hour peak of disease, which returned back to near-initial values by 48 hours. Additionally, *Asc^−/−^* mice displayed no significant differences in *S. anginosus* dissemination to the spleen or liver compared to wild type mice after 24 hours of infection, and this bacterial dissemination was also cleared at 48 hours post-infection in the *Asc^−/−^*tissue. Together, this indicates that the inflammasome promotes an acute and limiting proinflammatory response during *S. anginosus* infection, but it does not impact overall bacterial clearance. However, at 48 hrs post-infections, *Asc^−/−^* mice displayed elevated liver phospho-p65 compared to wild type controls, which may reflect an exaggerated inflammatory response resulting from loss of inflammasome-mediated detection of *S. anginosus* in the tissue of these mice. To determine which inflammasome sensor was involved in this acute response to *S. anginosus* infection *in vivo*, we infected *Aim2^−/−^*and *Nlrp3^−/−^* mice and tracked disease progression over 48 hours. *Nlrp3^−/−^* mice displayed a decrease in weight loss in comparison to wild type mice, similar to *Asc^−/−^* mice. This indicates that the NLRP3 inflammasome plays a role in innate immune response against *S. anginosus in vivo*, similar to what we observed during macrophage infection *in vitro*. No mortality was observed during *S. anginosus* infection of wild type or inflammasome-deficient animals, which was in stark contrast to the *in vivo* response to *S. pyogenes* when delivered via the same infectious route [37]. Interestingly, while infected *Nlrp3^−/−^* mice displayed similar morbidity as *Asc^−/−^*mice, *Aim2^−/−^* mice displayed more severe weight loss than wild type mice. Because *Aim2^−/−^*mice presented a phenotypically distinct response to *S. anginosus* infection compared to *Asc^−/−^* mice, these findings point towards an inflammasome-independent role for AIM2 during *S. anginosus* infection *in vivo*. This is in line with our findings that AIM2 was dispensable for *S. anginosus*-induced IL-1β production in macrophages. Indeed, AIM2 has inflammasome-distinct functions in non-immune cells (e.g., the epithelium) [34]. Additionally, we saw increased phospho-p65 in the spleen of *Aim2^−/−^* mice 48 hours after infection. This may be attributed to the noncanonical role of AIM2 as an inhibitor of the PI3K/AKT cell signaling pathway, which intersects with the NF-κB pathway [34], [49], [50]. However, the contribution of non-inflammasome functions of AIM2 during *S. anginosus* infection is outside of the scope of this study.

Although there are multiple ways *S. anginosus* can expand and/or become dysbiotic in the oral cavity, a major question that remains is how and why this species is elevated during OSCC. In normal epithelium, oral streptococcus such as *S. mitis*, *S. sanguinis* and *S. gordonii* are pioneer colonizers which form the structural architecture of the oral biofilm. This involves interactions with other streptococci species, promotion of microbial succession and fostering of diverse polymicrobial interactions in the oral cavity [51], [52]. These species restrict the colonization of other oral microbes through their production of alkali, bacteriocins, and hydrogen peroxide. Diet plays a large role in maintenance of early colonizing species and a high sugar diet along with poor oral hygiene can create an acidic environment that is less hospitable to these early commensal colonizers, which are not well-adapted to acidic conditions. This allows for the enrichment of acid-tolerant (aciduric) taxa such as *S. mutans* and favors cariogenic inflammatory environment leading to dysbiosis [53]. Additionally, the oral microbiome may become dysbiotic as a result of compromised immunity due to aging, as there is a positive correlation between *S. anginous* and age [54], much like OSCC. Other cancer-associated factors like smoking along with tissue injury or antibiotic use during infection can lead to dysbiosis of the oral microbiome as well [54]. While interspecies interactions of oral streptococci play an important role in limiting dysbiosis, the connection between *S. anginosus* and the immune system during localized and systemic inflammatory responses warrants more investigation, especially during the development and progression of cancer.

Our experiments largely explored the consequence of *S. anginosus*-macrophage interactions, but it is important to consider how these interactions would occur in healthy tissue versus oral pathologies like OSCC. While this is still an area of question for the oral cavity, *S. anginosus* infection of gastric tissue decreases expression of epithelial tight junction proteins claudin18, occluden and zonula occluden, leading to increased gastric epithelial permeability [16]. Additionally, *S. anginosus* increases expression of gastric chemokines CCL20 and CXCR2, which promote immune cell infiltration and increases both proinflammatory and immunosuppressive immune cell types, including Th17 CD4 T cells and tumor-associated macrophages, respectively [16]. *S. anginosus* is also associated with the recruitment of immune cells within human esophageal cancer as evidenced by an increase in chemo attractants IL-8 and GRO⍰ in esophageal tissue sections containing *S. anginosus* [55]. Overall, these findings highlight multiple mechanisms by which *S. anginosus* may promote interactions with immune cells like tissue-resident and recruited macrophages, but there is still much to learn about how *S. anginosus* interacts with different immune cells to impact proinflammatory responses. Here we show that *S. anginosus* replicates within macrophages, and its viability is essential to fully induce many downstream inflammatory pathways that require NF-κB, including COX2, iNOS, TNFα and inflammasome activation. The ability of *S. anginosus* to replicate intracellularly may be key to its observed sustained activation of the NF-κB pathway and inflammation within the epithelium and the tumor immune microenvironment. Our findings suggest *S. anginosus* may drive M1 differentiation of recruited and tumor associated macrophages (TAMs), which produce a variety of proinflammatory proteins and cytokines through the NF-κB pathway including COX2, iNOS, and TNFα. Increased COX2 and iNOS expression in TAMs may lead to increased VEGF and reactive oxygen species, respectively, which can exacerbate cancer by inducing angiogenesis and introducing new mutations through DNA oxidation [56]. TNFα is also increased in multiple cancer types and is associated with angiogenesis, tumor growth and DNA damage [57]. Both NF-κB and inflammasome activation are required for active IL-1β secretion, which is highly implicated during cancer [58]. Elevated IL-1β is present in a variety of cancers including OSCC [59], [60]. While acute production of inflammatory cytokines like IL-1β is essential to the clearance of many pathogens, prolonged IL-1β production and unchecked chronic inflammation is detrimental and may drive tumor progression through multiple mechanisms [35]. IL-1β may directly impact the epithelium to drive tumor growth and epithelial to mesenchymal transition through induction of MYC and downregulation of E-cadherin, respectively [61], [62]. Additionally, IL-1β can drive production of other pro-inflammatory molecules, which can cause cell death and release of DAMPs, which further contribute to disease progression [63]. Activation of the inflammasome for IL-1β secretion can occur by recognition of a variety of DAMPs and PAMPs. Here we found that *S. anginosus* is sensed by the NLRP3 inflammasome. NLRP3 and downstream IL-1β is largely implicated in several chronic inflammatory conditions and the development of OSCC [64]. Moreover, elevated NLRP3 in head and neck cancers including OSCC is associated with increased tumor invasion, survival and drug resistance [58]. However, the contribution of oral microbes like *S. anginosus* during NLRP3 overexpression and activation during cancer remains unclear. Dysbiosis and/or overgrowth of *S. anginosus* within the oral microbiome likely increases the presence of multiple damage- and pathogen-associated factors that trigger the NLRP3 inflammasome and could drive disease outcomes. As *S. anginosus* clearly impacts the TME in multiple ways, understanding its interactions with both epithelial cells and infiltrating immune cells will be key in understanding its impact on OSCC.

## Author Contributions

Conceptualization: A.M.A., S.S.K., and J.E.W. Methodology: A.M.A., D.M.R and J.E.W. Data Curation: A.M.A., C.S., S.S.K., and D.M.R. Formal Analysis: A.M.A., C.S., D.M.R., and J.E.W. Investigation: A.M.A, and J.E.W. Writing (original draft) A.A., S.S.K, and J.E.W. Writing (review and editing): A.M.A. and J.E.W. Resources: J.E.W. Supervision: J.E.W.

## Acknowledgements and Funding

This study was supported by an Arizona Biomedical Research Center New Investigator Award CTR056057 (J.E.W) and the University of Arizona Health Sciences Strategic Initiatives— Personalized Defense sanctioned under the Immune-Microbe Interface in Health and Disease (IMIHD) group (S.S.K). This work was also supported by NIH grants T32GM139779 (A.M.A), F31DE032263 (D.M.R), and T32AG058503 (D.M.R), along with a University of Arizona Cancer Center Cancer Biology Program Pilot Award.

We would like to thank Juanita L. Merchant at the University of Arizona Cancer Center for providing access to microscopy resources. We also thank the Experimental Mouse Shared Resource at the University of Arizona Cancer Center for technical assistance, which was supported by the National Cancer Institute of the National Institutes of Health under award number P30CA023074.

---

**Supplemental Figure 1. Tissue Western Blots from *S. anginosus*-infected WT and *Asc^−/−^*mice:** a) Western Blot of phospho-p65 and total p65 on protein homogenates from the spleen of WT and *Asc^−/−^* mice infected with 2×10^9^ CFUs of *S. anginosus* for 24 hours. b) Western Blot of phospho-p65 and total p65 on protein homogenates from the spleen of WT and *Asc^−/−^* mice infected with 2×10^9^ CFUs of *S. anginosus* for 48 hours. c) Western Blot of phospho-p65 and total p65 on protein homogenates from the liver of WT and *Asc^−/−^* mice infected with 2×10^9^ CFUs of *S. anginosus* for 48 hours. WT n=5, *Asc^−/−^* n=6 for 24-hour infections. WT n=4, *Asc^−/−^* n=4 for 48-hour infections.

**Supplemental Figure 2: Tissue Western Blots from *S. anginosus*-infected WT, *Aim2^−/−^* and *Nlrp3^−/−^* mice:** a) Western Blot of phospho-p65 and total p65 on protein homogenates from the spleen of WT, *Aim2^−/−^* and *Nlrp3^−/−^* mice infected with 2×10^9^ CFUs of *S. anginosus* for 24 hours. b) Western Blot of phospho-p65 and total p65 on protein homogenates from the liver of WT, *Aim2^−/−^* and *Nlrp3^−/−^* mice infected with 2×10^9^ CFUs of *S. anginosus* for 24 hours. c) Western Blot of phospho-p65 and total p65 on protein homogenates from the spleen of WT, *Aim2^−/−^* and *Nlrp3^−/−^* mice infected with 2×10^9^ CFUs of *S. anginosus* for 48 hours WT n=5, *Aim2^−/−^* n=5, *Nlrp3^−/−^* n=5 for 24-hour infections. WT n=5, *Aim2^−/−^* n=4, *Nlrp3^−/−^* n=5 for 48-hour infections.

