## Supplemental Figures 1+2 for "*Streptococcus anginosus* Activates the NLRP3 Inflammasome to Promote Inflammatory Responses from Macrophages"

### Slide 1
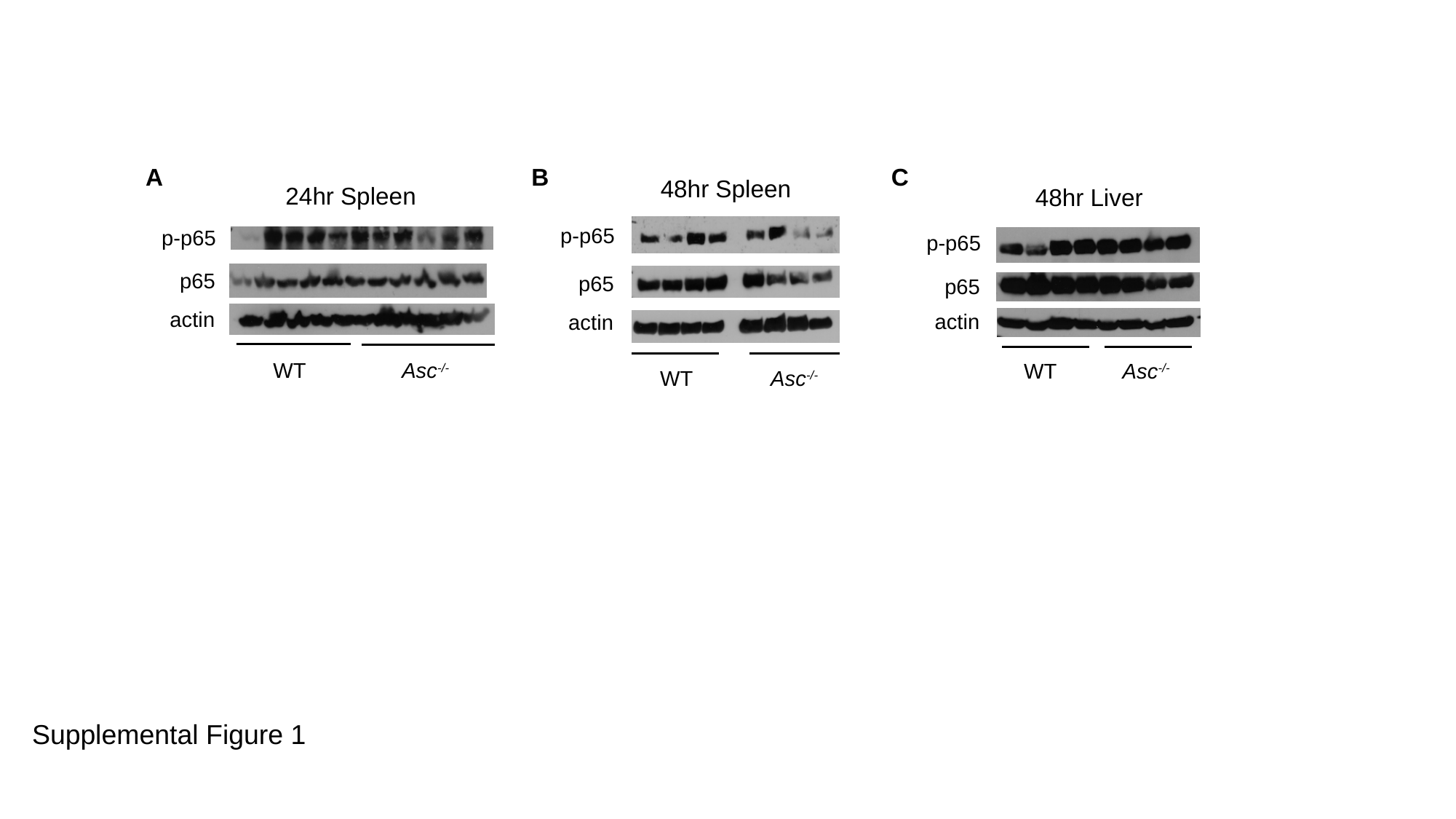

A B C
48hr Spleen
WT Asc^-/-^
p-p65
p65
actin
24hr Spleen
WT Asc^-/-^
p-p65
p65
actin
48hr Liver
p-p65
p65
actin
WT Asc^-/-^
Supplemental Figure 1

### Slide 2
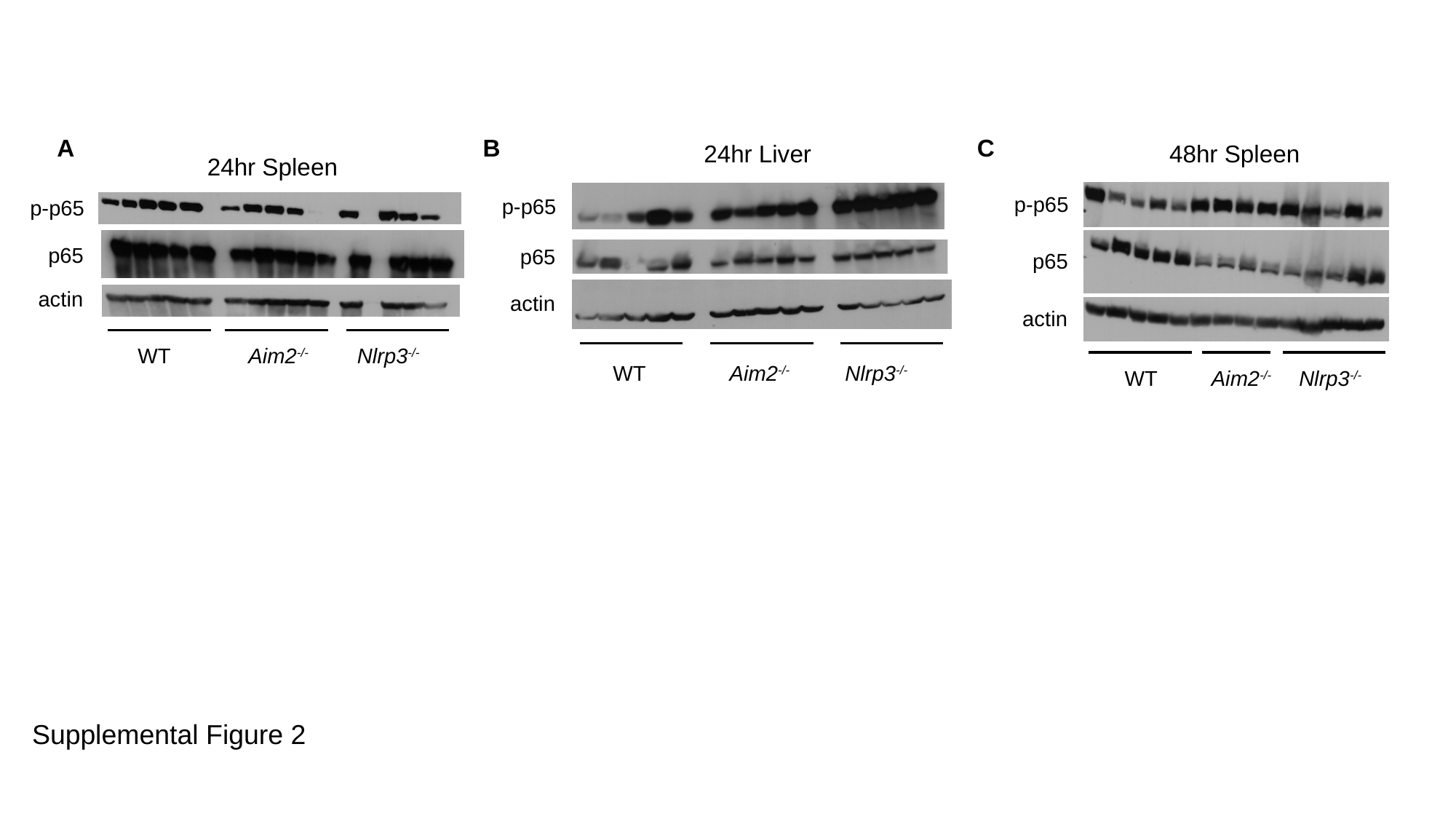

A B C
24hr Liver
p-p65
p65
actin
WT Aim2^-/-^ Nlrp3^-/-^
48hr Spleen
p-p65
p65
actin
WT Aim2^-/-^ Nlrp3^-/-^
24hr Spleen
WT Aim2^-/-^ Nlrp3^-/-^
p-p65
p65
actin
Supplemental Figure 2
